# A rapid approach for sex assignment by RAD-seq using a reference genome

**DOI:** 10.1101/2023.01.30.526394

**Authors:** Diego M. Peralta, Juan I. Túnez, Ulises E. Rodríguez Cruz, Santiago G. Ceballos

## Abstract

Sex identification is a common objective in molecular ecology. While many vertebrates display sexual dimorphism, determining the sex can be challenging in certain situations, such as species lacking clear sex-related phenotypic characteristics or in studies using non-invasive methods. In these cases, DNA analyses serve as valuable tools not only for sex determination but also for validating sex assignment based on phenotypic traits. In this study, we developed a bioinformatic framework for sex assignment using genomic data obtained through GBS, and having an available closely related genome assembled at the chromosome level. Our method consists of two *ad hoc* indexes that rely on the different properties of the mammalian heteromorphic sex chromosomes. For this purpose, we mapped RAD-seq loci to a reference genome and then obtained missingness and coverage depth values for the autosomes and X and Y chromosomes of each individual. Our methodology successfully determined the sex of 165 fur seals that had been phenotypically sexed in a previous study and 40 sea lions sampled in a non-invasive way. Additionally, we evaluated the accuracy of each index in sequences with varying average coverage depths, with Index Y proving greater reliability and robustness in assigning sex to individuals with low-depth coverage. We believe that the approach presented here can be extended to any animal taxa with known heteromorphic XY/ZW sex chromosome systems and that it can tolerate various qualities of GBS sequencing data.

## Introduction

Data on the genetic diversity, population size, sex ratio, kinship and distribution of free-ranging wildlife are essential for establishing effective conservation strategies [1]. In particular, information on the sex of individuals is of great importance in population ecology and conservation [2,3]. For example, it contributes to the understanding of population structure and dynamics in monitoring programs [4] or allows to identify and infer socio-ecological patterns in behavioral studies of living and extinct species [5,6], respectively. Knowledge of the sex ratio of effective individuals is particularly relevant for species or populations of conservation concern, as unbalanced sex ratios may have negative impacts on population growth and resilience [7]. However, sex identification is hampered in non-sexually dimorphic species (especially during the juvenile period) and in elusive ones due to the difficulty of direct observation [3,8]. As a result, the determination of sex through DNA analysis applied to biological samples (mainly those collected in a non-invasive way) has become a useful tool in molecular ecology and conservation genetics [9].

In mammals, molecular sexing has traditionally relied on the use of the polymerase chain reaction (PCR) for amplification of specific fragments on the Y chromosome or amplification of homologous fragments from both X and Y chromosomes [9–13]. Nevertheless, these methods depend on amplification reliability and external controls [14,15]. Other commonly used approaches are based on differences in size polymorphisms among X and Y homologous fragments such as Amelogenin gene size [16], or on the existence of unique restriction sites in a Y fragment such as zinc finger genes [17–19]. However, these methods do not apply to all taxa [15].

Restriction Associated DNA sequencing (RAD-seq) has emerged as a popular method in the field of population genetics because it overcomes the limitations of traditional techniques [20] and allows working with wild populations and non-model species, among other advantages. Since the advent of RAD-seq, several studies on sex assignment and sex chromosome systems, among others, have benefited from this technique. It improved the understanding of sex chromosomes systems, revealed the influence of sex-linked markers in population genetic studies, or gave tools to assign sex to samples from different sources with successful results [21,22]. For example, Carmichael *et al*. [23] identified a sex-linked SNP marker in a salmon’s parasite thus confirming the presence of a genetic mechanism for sex determination and pointed out that this information could be useful in the development of control strategies. Benestan *et al*. [24] stated that an unbalanced sex ratio in the samples and the presence of sex-linked markers (not removed from the dataset) may lead to a potential bias in the correct interpretation of population structure. In the context of individual sex identification, Stovall *et al*. [25] successfully assigned the sex to 86 New Zealand fur seals (*Arctocephalus forsteri*) through the discovery and statistical validation of sex-specific loci using a de novo RAD-seq approach. Their findings demonstrated that monomorphic loci (frequently ignored in the SNP datasets) are relevant when identifying sex-linked markers. Finally, Trenkel *et al*. [26] applied three sex-determining methods on 1680 rays of unknown sex-determination system and offered recommendations based on the error of each method. The main difference between the former two works and the third one is that it uses a reference genome.

In this context, the growing popularity of methods like RAD-seq [27] takes place simultaneously with the positive trend in generation of new reference genomes. In fact, the availability of genomes assembled at chromosome-level has shown a remarkable ten-fold increase from 2015 to the present day, mainly due to the coordinated and standardized efforts of international genome initiatives [28]. Taking mammals and birds as an example, there are 655 chromosome-level genome assemblies (of which 32% contain sex chromosomes) from 379 species that are currently available at NCBI (https://www.ncbi.nlm.nih.gov/datasets/genome/. Last accession 08-24-2023). Although there is overrepresentation within the clades of these groups (i.e., primates, artiodactyls, rodents and carnivores among mammals; and passeriformes, gallinaceous poultry and waterfowl among birds), this drawback is likely to be overcome by the continuous addition of new taxa.

In addition to the above, the number of genome assemblies containing sex chromosomes is continuously increasing [29], allowing for more comprehensive analyses of population and functional genomics. In particular, this contributed to an enhanced understanding of sex determination systems and their significant variability. In the animal kingdom, these are typically exemplified by the X/Y and Z/W configurations, in which one of the chromosomal pairs may potentially vary in size and carries sex-determining factors. The fundamental distinction between XY/ZW systems lies in the identity of the heterogametic sex (i.e., X/Z for males and Z/W for females) [29], even though various deviations have been reported. For instance, monomorphic forms of XY/ZW with chromosomal pairs being similar in shape and genetic content, as observed in some lizards [30]; a reduced number of sex chromosomes as observed in two subspecies of the Japanese rodent *Tokudaia osimensis* due to the lack of the Y chromosome [31]; or the presence of an additional X chromosome, as reported for the African pygmy mouse *Mus minutoides* [32]. Additionally, the spectrum encompasses a broad range of aberrations such as aneuploidies and neo-sex chromosomes, among others [29,33].

Considering the accuracy of RAD-seq and the increasing availability of reference genome assemblies including sex chromosomes in many species, here we propose a simple methodology for sexing individuals based on the different properties of the mammalian sex chromosomes X and Y and the possibility of mapping RAD-seq loci to a reference genome. Thus, we developed two indexes aiming at identifying the sex of animals with heteromorphic XY/ZW sex chromosomes, based on a closely related genome assembled at the chromosome level. The combination of both indexes allowed us to accurately assign sex to the fur seal dataset from Stovall *et al*. [25] and to the sea lion dataset from Peralta *et al*. [34].

## Methods

### Samples

In this study we used GBS (Genotyping-by-sequencing) data from 165 live fur seals (*Arctocephalus forsteri*) whose sex had been determined phenotypically (93 females and 72 males). These were obtained from a GBS dataset of 255 dead and live individuals [25] available at NCBI Sequence Read Archive (BioProject Accession no.: PRJNA419445). Then, we applied the method to the RAD-seq dataset of 40 sea lions (*Otaria flavescens*) of unknown sex from Peralta *et al*. [34] available at NCBI Sequence Read Archive (BioProject Accession no.: PRJNA1022835). Samples were collected by either invasive or non-invasive methods from individuals along the coast of Argentina [35].

### RAD sequencing and filtering

The RAD-seq library from Peralta *et al*. [34] was constructed according to the protocol described in Roesti *et al*. [36,37], adopted from Hohenlohe *et al*. [38]. The restriction digestion was made using only the Sbf1 enzyme, followed by the addition of a specific barcode to the restricted DNA. The library was single-end sequenced to 150 bp in a single lane on an Illumina HiSeq 4000 at the Genomics and Cell Characterization Core Facility, University of Oregon, USA.

Individual reads of both data sets were mapped to the California sea lion (*Zalophus californianus*) representative genome mZalCal1.pri.v2 (GCA_009762305.2) using BWA MEM v0.7.17 [39] with default parameters. Unmapped reads were removed from the SAM alignment files using SAMtools v1.7 [40]. For detailed methodology see Peralta *et al*. [34]. Regarding the selection of the representative genome, it is worth mentioning that the Pinnipedia superfamily is characterized by a pronounced karyotypic uniformity, particularly within Otariidae, having 2n = 36 and closely agreeing karyotypes [41]. This allowed us to use the *Z. californianus* genome as representative of both species.

The filtering steps (gstacks, STACKS v2.64) [42,43] were aimed to retain as much sex-linked markers as possible. Therefore, considering that Y-linked loci are present only in males, we set the minimum proportion of individuals across populations required to process a locus (-R) to 0.3, which assumes a minimum of 30% for males in our dataset. We also filtered loci with a maximum observed heterozygosity greater than 0.7 (--max-obs-het), and used the flag to create a vcf file containing all sites (variable and fixed) within the RAD loci (--vcf-all).

### Sex identification

Sex identification of individuals was carried out using the chromosome-level assembly of the California sea lion genome as reference. Since male mammals carry a single copy of the X chromosome, they are expected to have about 50% of coverage depth for loci mapping to the X chromosomes compared to an autosomal chromosome, while in females this value should be similar when compare the X and any autosome. In regard to the Y chromosome, females are expected to have a high value of missingness (defined as the percentage of total number of loci found in the population that are missing in an individual), while in males it should be similar to the missingness of any other chromosome. Theoretically, however, it should be more similar to that of the X chromosome because the heterogametic sex has only a single copy of each sex chromosome. Taking all this into account we constructed two different indexes for sex identification. We defined Index X as the average coverage depth of the X chromosome divided by the average coverage depth of the autosomes. On the other hand, to create Index Y, we first defined the individual completeness as one minus missingness and used the completeness values to get an index with expected values ranging between 0 and 1 (See Table 1). Index Y could be interpreted as the expected missingness for the Y chromosome in relation to the overall missingness in that individual.

**Table 1.** Index X and Y calculation.

|  |  | Expected Value |  |
| --- | --- | --- | --- |
|  | Equation | Female | Male |
| Index X | $= \frac{Depth_X}{Depth_A}$ | 1 | 0.5 |
| Index Y | $= \frac{Com_X - Com_Y}{Com_X}$ | 1 | 0 |

To calculate missingness and coverage depth we used different options of VCFtools [44] over the output files of STACKS “populations.all.vcf” [42,43]. We filtered RAD-loci in the Y (NC_045613.1) and X (NC_045612.1) chromosomes, and all the autosomes to obtain the missingness per individual with the option “--missing-indv” and the average coverage depth with the option “--depth” in VCFtools. We used all the positions in the RAD loci because the indexes do not depend on the presence or absence of SNPs in the loci but rather on the presence or absence of the loci themselves. A summary of the step-by-step process is depicted in the workflow in Fig 1, while the detailed procedure is available in the GitHub repository at https://github.com/Dieggarp/sexing (S1 File). Next, we calculated both indexes with the equations given in Table 1 using R [45]. This last step was executed using an automated R script, “sexing.R” (S1 File), also included in the GitHub repository. This script creates two files, first “final_sexing.csv”, which contains the assignment of sex for each individual and second “sexing_plots.pdf”, which contains three dispersion plots (Index X vs Y, Index X vs coverage depth, Index Y vs coverage depth) to visualize the results. We only used values of Index Y for assigning sex to each individual because Index X may lead to sex misidentification (see Discussion).

**Fig 1.**
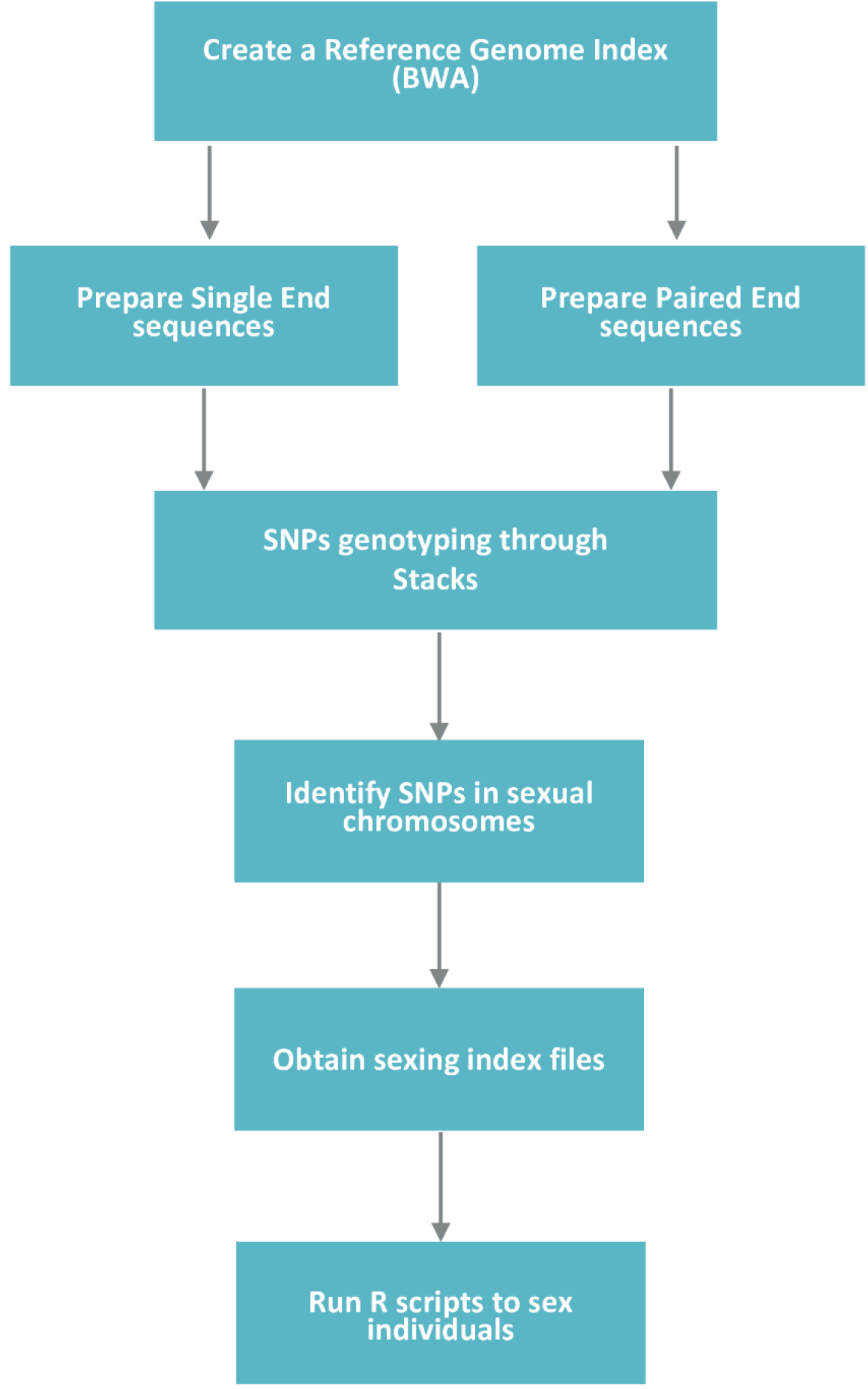
Sexing workflow. Illustration of a step-by-step summary of the entire sexing procedure, starting with the alignment to a reference genome and ending with the creation of two files that indicate the sex assigned to each individual.

Indexes X and Y for sex identification. Index X is computed by dividing the average coverage depth of the X chromosome by the average coverage depth of the autosomes, expecting to find equal values of average coverage depth for the X chromosome in females, obtaining an Index X = 1, and half in males, Index X = 0.5. Index Y is computed by taking the difference in completeness between the X and Y chromosomes and dividing it by the completeness of the X chromosome. As a result, females would exhibit a complete difference in completeness between the sex chromosomes, obtaining an Index Y = 1, given the absence of the Y chromosome (Y missingness = 1). In contrast, males would exhibit no difference in completeness between these chromosomes, resulting in Index Y = 0 due to the presence of both sex chromosomes.

DepthX and DepthA: average coverage depth across loci for the X chromosome and autosomes, respectively. ComX and ComY: completeness for the X and Y chromosomes, respectively. Com = one minus missingness, where missingness is the proportion of total number of loci in the population that are absent in that individual.

## Results

### RAD loci filtering

After quality and coverage filtering the *A. forsteri* dataset, we obtained an average of 2.9 million reads per sample. A total of 560,778 widely shared loci were recovered with an average coverage depth of 11.5 X ± 7.1. On average, 31,775.1±8,096.47 loci were found on autosomal chromosomes, 19,574 on the X chromosome and 551 on the Y chromosome. The former filtering with population parameters using “-r” of 0.3 resulted in 180,343 shared loci (present in at least 30% of samples), containing 16,690,597 sites (fixed and variable) used for subsequent analysis.

For the *O. flavescens* dataset we obtained an average of 3.2 million reads per sample. A total of 818,244 widely shared loci were recovered with an average coverage depth of 13.8 X ± 13.1. On average, 46,414.8±12,839.35 loci were found on autosomal chromosomes, 26,457 on the X chromosome and 1,169 on the Y chromosome. The former filtering with population parameters applying “-r” of 0.3 resulted in 127,983 shared loci (present in at least 30% of samples) containing 18,438,603 sites (fixed and variable), which were used for subsequent analysis.

### Genomic sexing

Of the 165 fur seals from Stovall *et al*. [25], we successfully sexed 164 (99.4%) individuals with both indexes and one with Index Y only, resulting in 95 females and 70 males (vs. 93 females and 72 males that had been phenotypically identified) (S1 Table). Interestingly, our two indexes sexed as females two individuals phenotypically classified as males [25]. Thus, there was 98.8% of coincidence between our method and sex assignment by phenotypic traits (but see discussion), 100% for females and 97.2% for males.

The dispersion plot for both indexes (Fig 2) showed two tight clouds of points in accordance with what was expected for males and females. On the one hand, tightly clustered points at the intersection of value 1 for both indexes were inferred to belong to females. On the other hand, points located almost entirely near the intersection of value 0.65 for Index X and around 0 for Index Y were inferred to belong to males. The value of the outlier point lying within the lower cloud of points was lower than the others (−1.6) for Index Y and close to 1 for Index X. It is worth noting that this individual also showed the lowest average coverage depth (1.4X), allowing us to test how this value affected both indexes. We found that low values of average coverage depth led to a remarkable decline in the performance of Index X (i.e., an individual of low average coverage depth was misidentified) but not of Index Y (S1 Fig). It is also interesting to remark that in male individuals, samples with very low average coverage depth can even give negative values for Index Y, thus improving sex identification.

**Figure 2.**
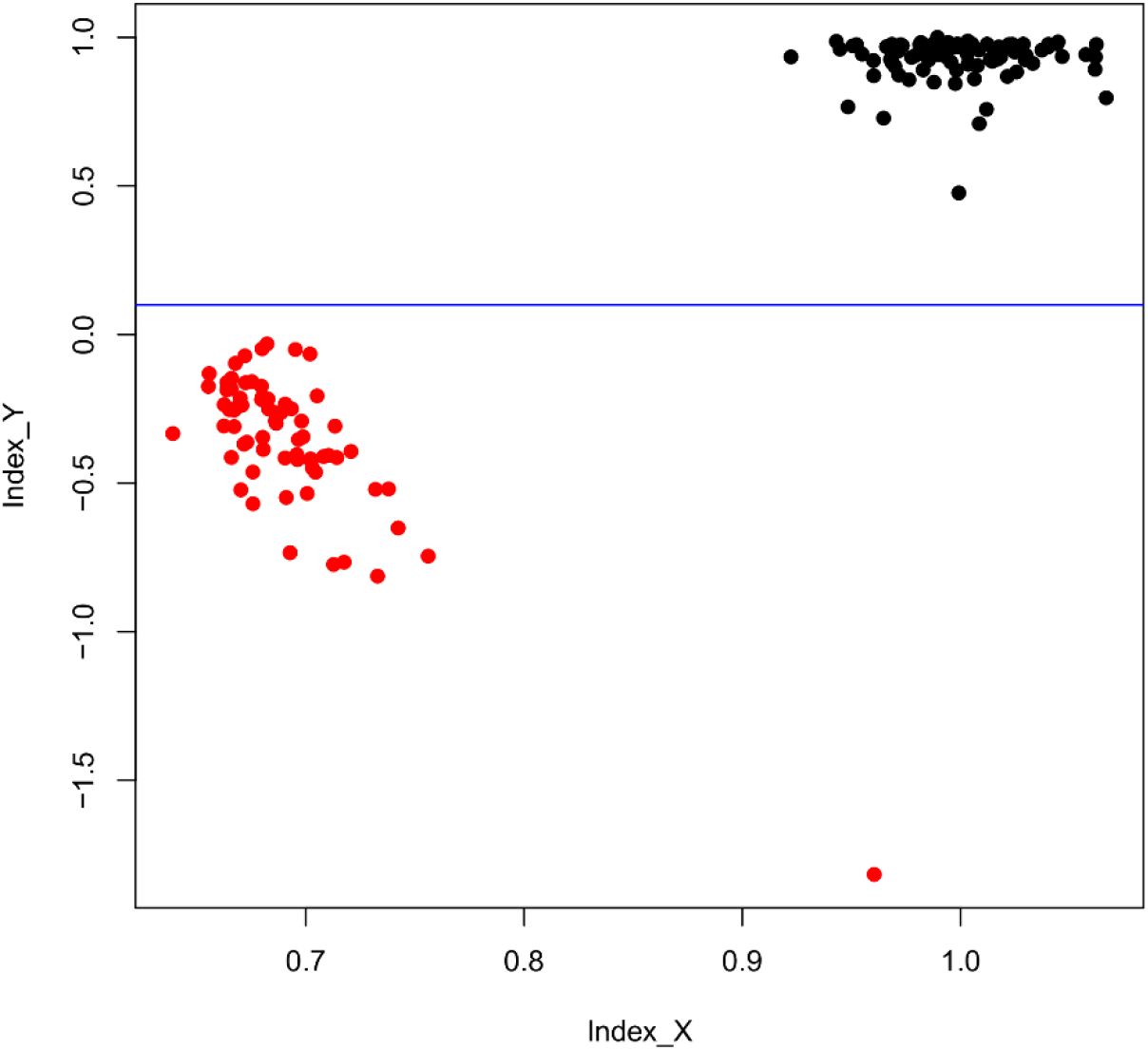
Dispersion plot of Index Y and X for fur seals. Dispersion plot of both indexes based on the dataset from Stovall *et al*. [25]. Red points: males; black points: females. The blue line indicates that Index Y achieves a complete separation between sexes.

Of the 40 individuals analyzed from the South American sea lions, 32 were successfully sexed using both indexes, resulting in 19 males and 13 females. The remaining individuals could only be sexed by Index Y, resulting in two females and six males. (S2 Table).

The dispersion plot for the X and Y indexes (Fig 3) showed two clouds of points representing males and females. The cloud of points corresponding to females was clustered at the intersection of value 1 for both X and Y, while that of males was near the intersection of values 0.5 for Index X and 0 for Index Y. However, some values were above 0.5 for Index X and below 0 for Index Y. Once more we evaluated the effect of coverage depth on our indexes, finding that Index X was the only one affected (S2 Fig) and that again male samples with very low average coverage depth (<10X) yielded negative values for Index Y, providing a better separation of sexes.

**Figure 3.**
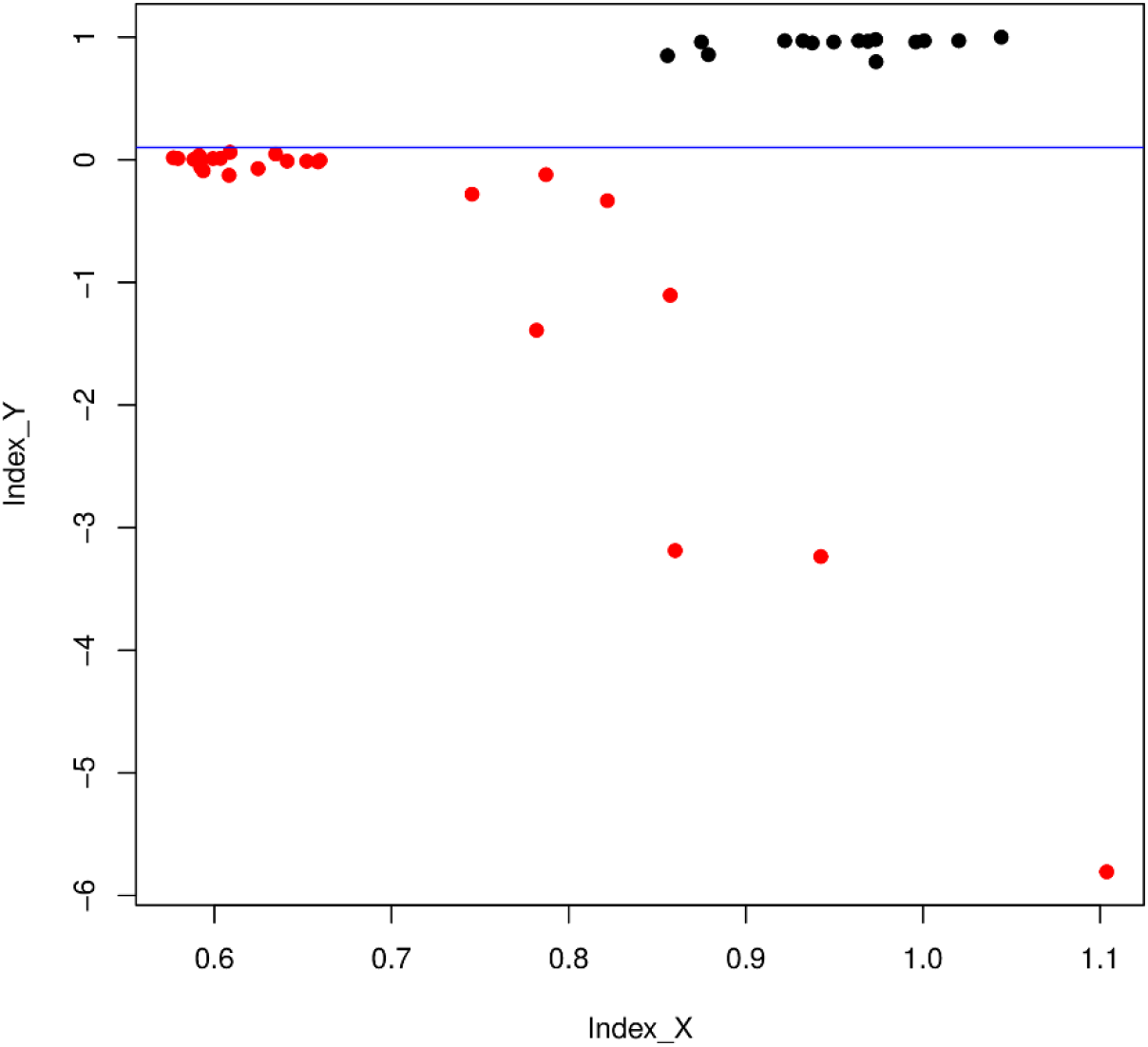
Dispersion plot of Index Y and X for sea lions. Dispersion plot of both indexes based on the dataset from Peralta *et al*. [34]. Red dots: males; black points: females. The blue line indicates that Index Y achieves a complete separation between sexes.

## Discussion

### Sexing animals from a RADseq approach

Here we provide a simple framework for sexing different types of samples using two *ad hoc* indexes. Our method will allow researchers working with RAD-seq or other similar GBS methodology and having an available reference genome, to sex individuals in an easy and reliable way.

After applying our indexes to the individuals in the Stovall *et al*. [25] dataset, we obtained 95 females and 70 males. The dispersion plot constructed with the indexes (Fig 2) showed that Index Y located females near the value 1 and males below 0. Index X grouped almost all individuals into two different clouds of points along the abscissa, except for one individual that could be differentiated only by the Index Y. It is important to highlight that our indexes identified as females two individuals previously labeled as males. In fact, Stovall *et al*. [25] reported that three individuals phenotypically labeled as males lacked male-specific genetic markers at the time of genotypic identification, thereby assuming that their sex had been misidentified. These cases of misidentifications or mislabeled data were further confirmed by one of the authors (Neil J. Gemmell, pers. comm.).

Our sea lion dataset yielded 25 males and 15 females, but eight of them could only be identified with Index Y because Index X could not discriminate between sexes. Certainly, the dispersion plot (Fig 3) showed that Index Y proved to be successful for sex determination while Index X failed in several cases. These results support the notion that Index Y is more reliable and robust than Index X.

The samples that could only be sexed by Index Y shared low levels of average coverage depth, with values less than 5X in some cases. This may be linked to the origin of the samples and the extent of DNA degradation, as indicated in Supplementary Table 2. Specifically, only one of the samples from the dataset of Peralta *et al*. [34] was obtained from a live animal while the remaining ones corresponded to deceased animals at different stages of decomposition. In this regard, we found that Index X requires an average coverage depth above 10X to correctly assign the sex of samples; below this value it tended to fail, with a bias toward males. A possible explanation may be that the difference in coverage depth between the X chromosome and the autosomes that characterizes males is reduced when average coverage decreases. On the contrary, the Y index seems to overcome this problem, as low coverage depth would not interfere with correct sex assignment. Index Y even seems to benefit from low coverage depth because males tend to show more negative values at lower coverage, resulting in a more pronounced distinction between sexes. In this sense, the influence of the different coverage depths on our indexes was evident in both the fur seal and our sea lion datasets. In turn, these inferences become more robust when contrasting the phenotypic sex of the seals with the efficacy of each of our indexes, confirming our suspicions about their different performance.

Overall, we suggest that Index Y is more reliable in assigning sex to individuals with low depth coverage, which is of particular relevance for studies involving non-invasive samples where DNA is usually degraded and on which RAD-seq has a weaker performance [46]. However, its joint use with Index X may provide a more accurate sexing result or can serve as a double check.

Our Index X is similar to the approach developed by Pečnerová *et al*. [5], who determined the sex of 98 woolly mammoths from frozen bone samples. They mapped the entire genome of mammoths against the chromosome-level LoxAfr4 assembly of a female African savannah elephant to compare the number of reads mapping to an autosome and to chromosome X. Like in our method, males were expected to have about 50% of reads mapping to chromosome X compared to the autosome, while females were expected to have a comparable number of reads mapping to both chromosomes. The main difference with the present study is that we developed Index X using a RAD-seq approach and compared values of coverage depth for all the sites (fixed and variable) in the RAD-loci identified, while Pečnerová *et al*. [5] used the number of total reads mapping to chromosome 8 and X, respectively, for each individual. Moreover, the method described by these authors requires the chromosomes under comparison to be of the similar length because it uses the number of total reads, while ours does not because it measures average coverage depth per loci.

Another method that deserves mention was used by Stovall *et al*. [25]. They employed *a de novo* approach to identify SNPs representing a useful tool if no chromosome-level reference genome is available for the species under study. However, multiple statistical validations must be performed to identify sex-related SNPs and assign them to individuals; this procedure needs to be repeated for each new study, thereby hampering its implementation. Moreover, it may be influenced by sequencing standards such as coverage depth and sample size. In comparison, the method proposed here is robust, accurate and easy to use even with samples having low average coverage depth and can be applied to new species with minimal modifications. Lastly, the accuracy of their sexing method compared to phenotypic sexing was 98.9% and 95.8% for females and males, respectively, while in the present study it was 100% and 97.2% for females and males, respectively.

Sex-associated markers have been used in studies covering a wide spectrum of topics such as sex-associated SNPs and sex-determining systems in different taxa including mammals, fishes, cnidarians, frogs, lobsters, magnolias and cannabis, among others [21,23–26,47–51]. In this context, it is worth mentioning the bioinformatic tool “RADsex”, which is used to identify sex-associated markers through statistical analysis and prediction models using RAD-seq data [21]. Among other benefits, this method also provides the possibility to assign sex to individuals based on genomic information.

Finally, the method described here is simple and useful for determining the sex of samples from different sources, but it can only be applied in species with heteromorphic sex chromosomes (e.g., XY/ZW). In these cases, one sex is heterogametic and sex chromosomes differ from each other in their features, such as size, gene content, repeat content, or structural differences. These characteristics give meaning to our indices and enable the detection of differences between males and females, achieving effective sex assignment.

The availability of a chromosome-level reference genome from a phylogenetically closely-related species may be regarded as another limitation of our method. However, reference genomes with these characteristics are progressively becoming more prevalent for non-model species, due to the progressive decrease in costs and DNA input requirements for genome assemblies, advancements in bioinformatics tools and improvements in annotation pipelines [52–55]. Additionally, ongoing advances in assembly technologies are addressing the challenges associated with assembling sex chromosomes [29,55].

All the above mentioned, along with the existence of numerous international consortia and genome projects focused on the creation of high-quality reference genomes for species across the phylogenetic tree of life (e.g., the 10,000 Plants Genome Sequencing Project, Earth BioGenome Project, Global Invertebrates Genomics Alliance, Vertebrate Genome Project), promise the publication of thousands of genome assemblies containing sex chromosomes within the next decade [29].

## Conclusion

We developed a simple methodology for determining the sex of different individuals using genomic RAD-seq data. Our approach comprises two *ad hoc* indices that successfully assigned the sex of sea lions and fur seals. Genomic libraries were created using DNA obtained from various sample types, including non-invasive ones, which sometimes hamper sex identification. We believe that this method could be extrapolated to any animal species with known heteromorphic XY/ZW sex chromosome systems, for which a chromosome-level reference genome of a closely-related species is available.

## Acknowledgments

We are grateful to the following people and organizations for their assistance during sampling: the park rangers of the National Parks and Protected Areas; the government agencies and their staff (Administración de Parques Nacionales, Secretaría de Medio Ambiente y Desarrollo Sustentable de Río Negro, Secretaría de Ambiente de Santa Cruz, Secretaría de Ambiente, Desarrollo Sostenible y Cambio Climático de Tierra del Fuego); the staff of the NGO Fundación Cadace; Adolfo Imbert, Dario Urruti and Agustín “Chepan” Szczepañsk from Centro Hípico del Fin del Mundo, Ushuaia; Lida Pimer; and Proyecto IMMA. We wish to thank Prof. Neil Gemmell for their time and willingness to answer our questions and the anonymous reviewer for manuscript feedback.

## Data availability

All relevant data are within the manuscript and its Supporting Information files. Moreover a detailed sequential overview of the methodological procedure is available at: https://github.com/Dieggarp/sexing. RAD-seq sequence reads belonging to Peralta et al. [34] are deposited in GenBank under the BioProject Accession no.: PRJNA1022835.

## Supporting information

**Supplementary File S1:** Step-by-step protocol. It will be also available on protocols.io.

**Supplementary figures:** Contains S1 Fig. and S2 Fig. with the dispersion plots of Indexes Y and X vs average coverage depth of both datasets.

**S1 Table:** Table of the parameters and index calculation for the fur seal (*Arctocephalus forsteri*).

**S2 Table:** Table of the parameters and index calculation for the sea lion (*Otaria flavescens*).

